# Modulation of temporal prediction by STN-DBS in Parkinson’s disease: Links between behavior and cortical oscillations

**DOI:** 10.1101/2025.05.29.656824

**Authors:** Rebecca Burke, Marleen J. Schoenfeld, Christian K.E. Moll, Alessandro Gulberti, Monika Pötter-Nerger, Andreas K. Engel

## Abstract

**BACKGROUND:** Accurate temporal prediction is essential for adaptive behavior and relies on coordinated neural activity within cortico-basal ganglia circuits. Parkinson’s disease (PD), with the main hallmark of dopaminergic depletion and abnormal neural synchrony, impairs this ability. Deep brain stimulation deep brain stimulation (DBS) of the subthalamic nucleus (STN) is a widely used treatment for reducing motor symptoms in PD, but its effects on temporal prediction remain not fully understood.

**OBJECTIVES:** This study aimed to investigate how STN-DBS influences temporal prediction performance and its underlying oscillatory dynamics in PD patients, with a particular focus on beta-band power and delta-band inter-trial phase consistency (ITPC).

**METHODS:** 13 PD patients (5 female, age: 64 ± 5.7 years; disease duration: 11.8 ± 1.8 years) with STN-DBS performed a temporal prediction task with (DBS ON) and without (DBS OFF) stimulation, while 64-channel-electroencephalography (EEG) was recorded. 20 age-matched healthy controls completed the same task. Behavioral performance was assessed using psychometric function slopes. Time-frequency analyses and source-level EEG measures examined beta power and delta ITPC.

**RESULTS:** PD patients showed impaired temporal prediction performance compared to controls, reflected in shallower psychometric slopes. DBS significantly improved performance to a level comparable to those of controls. EEG revealed reduced beta suppression in PD patients during DBS OFF, while beta suppression in DBS ON was comparable to controls. Both DBS OFF and ON exhibited reduced delta ITPC compared to controls. In DBS ON, source-level delta ITPC was positively correlated with temporal prediction accuracy.

**CONCLUSION:** STN-DBS improves temporal prediction performance in PD, likely through modulation of beta and delta oscillatory activity. While beta power suppression is partially restored, deficits in delta phase alignment persist, suggesting frequency-specific DBS effects on temporal prediction processes.

## INTRODUCTION

Accurate perception and prediction of time are fundamental for adaptive behavior, enabling humans to anticipate and respond to events in their environment. This ability involves estimating durations in the seconds-to-minutes [Coull and Nobre, 1998, Meck, 1996] and sub-seconds range [Smith et al., 2007] and is central to cognitive and motor processes. A key neural substrate underlying interval timing is the basal ganglia, a complex network of structures involved in motor, cognitive, and associative functions. Parkinson’s disease (PD), a neurodegenerative disorder characterized primarily by the progressive loss of dopaminergic neurons in the substantia nigra pars compacta, frequently disrupts these functions [Brown, 2003, Oswal et al., 2013]. The resulting dopamine deficiency slows the pace of the internal clock, leading to impairments in time perception and interval timing tasks [Meck, 1996, Rammsayer and Classen, 1997, Perbal et al., 2005]. Dopaminergic medication has been shown to ameliorate these deficits by modulating the internal clock and enhancing attentional control of temporal information [Jones et al., 2008]. However, despite medication, timing performance in PD remains heterogeneous, suggesting the involvement of additional neural mechanisms [Merchant et al., 2008]. Next to standard dopaminergic medication, deep brain stimulation (DBS) of the subthalamic nucleus (STN) is an established treatment for PD, primarily used to alleviate motor symptoms by reducing excessive beta synchronization within the cortico-basal ganglia loops [Oswal et al., 2013]. Beyond motor improvements, DBS has been shown to have heterogeneous effects on non-motor functions, including executive processes, working memory, and cognitive flexibility [Jahanshahi et al., 2000, Witt et al., 2004, Gülke et al., 2022, Yamanaka et al., 2012]. Specifically, DBS appears to mitigate PD-associated impairments in time interval memory retrieval [Wojtecki et al., 2011]. By modulating STN activity, DBS provides a unique opportunity to investigate the contribution of the basal ganglia in temporal prediction. In this study, we sought to examine how anticipatory neural dynamics associated with temporal prediction are influenced by DBS in PD. Using electroencephalography (EEG) and a well-established temporal prediction task [Burke et al., 2025, Daume et al., 2021], we compared oscillatory markers of anticipation in PD patients in one session with bilateral therapeutic STN-DBS turned on (DBS ON), and in another session with the device switched off (DBS OFF) to those in healthy age-matched controls. Our findings aim to bridge gaps in understanding the role of the basal ganglia in temporal processing and extend previous work on the cognitive effects of DBS [Jahanshahi et al., 2000, Pillon et al., 2000]. We expected that individuals with PD would show impairments in temporal prediction relative to age-matched healthy controls, reflected in greater variability in their timing judgments. This deficit was expected to appear as a shallower slope in psychometric function. Additionally, we hypothesized that these behavioral differences between patients and healthy controls can be linked to altered beta-band modulation, in line with previous findings [Gulberti et al., 2015, 2024], as well as reduced phase consistency of delta oscillations in the parietal and frontal cortices following stimulus offset. We further presumed that DBS might influence temporal judgments in PD patients, resulting in systematic changes in the psychometric function, either by shifting the perceived timing or by modifying the slope.

## MATERIALS AND METHODS

### Participants

13 patients (5 female, mean age: 64 *±* 5.7 years) with a diagnosis of advanced idiopathic PD (mean disease duration: 11.8 *±* 1.8 years) and 20 healthy age-matched controls (13 female, mean age: 60.3 *±* 3.9 years) took part in the study. The experiment was conducted in accordance with the Declaration of Helsinki and approved by the ethics committee of the Hamburg Medical Association. All participants gave written informed consent and received monetary compensation. Handedness was assessed using the short version of the Edinburgh Handedness Inventory [Oldfield, 1971]. All participants reported normal or corrected-to-normal vision and no history of psychiatric diseases. Participants were excluded if they showed indications of dementia based on a global dementia screening battery (Mini-Mental State Exam score <25) or psychiatric diseases (per DSM-IV) or neurological conditions other than PD or were receiving medication for depression. PD patients underwent therapeutical bilateral implantation of DBS electrodes in the STN prior to the experiment (months since surgery: 23.1 *±* 15.4) and their levodopa-equivalent daily dose (LEDD, conversion factors as described by Jost et al. [2023]) was 786.6 *±* 547.4 mg at the time of participation (see Table 1). To demonstrate patients’ disease progression and severity, we derived the motor-subscore (III) of the MDS-Unified Parkinson’s Disease Rating Scale (UPDRS) [Goetz et al., 2008] and the Hoehn & Yahr scale (H&Y) [Hoehn and Yahr, 1967] from patients’ medical history. Controls reported no history of neurological or psychiatric disorders. Both patients (for both sessions in DBS OFF and ON) and controls completed the Trail Making Test A and B [Reitan, 1956], as well as the digital version of Berg’s Card Sorting Test [Grant and Berg, 1948], to assess their executive functions, specifically cognitive flexibility and logical reasoning. Patients and controls were excluded if they could not perform these tests. Since our focus in the present study was on effects of STN-DBS on temporal prediction, all patients were on their regular medication for all recordings. This also ensured patients’ physical comfort during the measurement. While the present manuscript focuses on the comparison between PD patients and age-matched healthy controls, the control data will also be included in a separate publication examining age-related differences between younger and older adults.

**Table 1:** Clinical and demographic characteristics of PD patients. In column “DBS parameters” values reported are: stimulation frequency in Hz, active contacts, impulse amplitude in volts and impulse width in *µ*s for the left and right electrode, respectively. For the left Medtronic (ME) electrode, contact 0 was the most ventral and contact 3 was the most dorsal. For the right ME electrode, contact 8 was the most ventral and contact 11 was the most dorsal. For the left Boston Scientific (BS) electrode, contact 1 was the most ventral and contact 8 was the most dorsal. For the right BS electrode, contact 9 was the most ventral and contact 16 was the most dorsal. Abbreviations: H&Y = Hoehn & Yahr scale. Op = operation for DBS electrodes implantation. UPDRS-III = Unified Parkinson’s Disease Rating Scale of the Movement Disorder Society, motor-subscore (part III). LEDD = levodopa equivalent daily doses.

| Case<br>Gender<br>Age | Disease<br>duration<br>(years) | Pre-op<br>H&Y | Pre-op<br>medication<br>(LEDD) | Pre-op UPDRS |  | DBS<br>device | DBS parameters for:<br>Left electrode (1. row)<br>Right electrode (2. row) | Post-op<br>H&Y | Post-op<br>medication<br>(LEDD) | Post-op UPDRS |  |
| --- | --- | --- | --- | --- | --- | --- | --- | --- | --- | --- | --- |
|  |  |  |  | DOPA-OFF | DOPA-ON |  |  |  |  | DOPA-OFF<br>DBS-ON | DOPA-ON<br>DBS-ON |
| 1, M, 72 | 5 | 2 | 300mg | 12 | 21 | ME | 130Hz, 1-, 2.0V, 60μs<br>130Hz, 9-, 2.0V, 60 μs | 2 | 100mg | 20 | 26 |
| 2, M, 65 | 12 | 2 | 1646mg | 11 | 42 | ME | 130Hz, 1-, 2-, 6.0V, 60μs<br>130Hz, 9-, 5.5V, 60μs | 2 | 713mg | 13 | 19 |
| 3, F, 59 | 12 | 3 | 1413mg | 13 | 29 | BS | 130Hz, 2-, 1.5V, 60μs<br>130Hz, 2-, 1.7V, 60μs | 1 | 963mg | 5 | 6 |
| 4, F, 63 | 11 | 2 | 1449mg | 13 | 21 | ME | 140Hz, 1-, 7.5V, 60μs<br>140Hz, 9-, 6.6V, 60μs | 2 | 255mg | 10 | 15 |
| 5, M, 60 | 10 | 2 | 1875mg | 17 | 23 | BS | 130Hz, 2-, 1.1V, 60μs<br>130Hz, 2-, 2.0V, 60μs | 2 | 630mg | 11 | 12 |
| 6, M, 72 | 15 | 2 | 1150mg | 17 | 41 | BS | 130Hz, 1-, 4.3V, 60μs<br>130Hz, 1-, 5.7V, 60μs | 2 | 1253mg | 12 | 33 |
| 7, M, 58 | 11 | 2 | 924mg | 9 | 18 | BS | 130Hz, 2-, 1.8V, 60μs<br>130Hz, 2-, 1.4V, 60μs | 1 | 1862mg | 7 | 15 |
| 8, F, 58 | 10 | 2.5 | 1121mg | 9 | 24 | BS | 128Hz, 1-, 2-, 6.7V, 60μs<br>128Hz, 2-, 1.4V, 60μs | 2 | 350mg | 12 | 16 |
| 9, M, 66 | 11 | 1 | 1442mg | 14 | 24 | BS | 130Hz, 2-, 4.8V, 80μs<br>135Hz, 2-, 4.5V, 80μs | 2 | 1692mg | 9 | 17 |
| 10, F, 71 | 13 | 2 | 1325mg | 11 | 35 | BS | 130Hz, 3-, 2.5V, 60μs<br>130Hz, 2-, 2.3V, 60μs | 0 | 507mg | 9 | 25 |
| 11, M, 70 | 15 | 2.5 | 1181mg | 17 | 22 | BS | 130Hz, 3-, 2.0V, 60μs<br>130Hz, 3-, 2.3V, 60μs | 2 | 971mg | 11 | 20 |
| 12, F, 63 | 10 | 3 | 1000mg | 12 | 46 | BS | 130Hz, 3-, 4.0V, 60μs<br>130Hz, 2-, 4-, 1.7V, 60μs | 2 | 405mg | 19 | 34 |
| 13, M, 59 | 12 | 3 | 1782mg | 11 | 31 | ME | 125Hz, 1-, 5.4V, 60μs<br>125Hz, 9-, 3.9V, 60μs | 2 | 525mg | 5 | 9 |

### Experimental Paradigm

The study was conducted in two sessions (two different recording days within one week), maintaining consistent timing of the day and response mapping. In each session, participants performed the neuropsychological tests followed by the temporal prediction task (see Figure 1A; for details on experimental parameters see Daume et al. [2021] and Burke et al. [2025]. Participants were seated in a dimly lit chamber that was both electrically shielded, and sound attenuated. Visual stimuli were presented on a matte LCD screen (1920 × 1080 px, 120 Hz refresh rate). Throughout the experiment, participants were asked to fixate a red fixation dot surrounded by a white noise occluder. Each trial began with a fixation period lasting 1500 ms. Following this, a white ellipse appeared in the left periphery of the screen and moved towards the occluder at a constant velocity. The starting position of the ellipse varied, resulting in movement intervals lasting between 1000 and 1500 ms. After a delay *t*, it reemerged on the right side and continued moving for 500 ms. Reappearance times varied randomly by Δ*t* (*±*34 to *±*934 ms, in 100 ms steps), relative to the objectively correct 1500 ms interval. Participants judged whether the reappearance was *too early* or *too late* via right-hand button press, based on motion perception. Response-button mapping was counterbalanced across participants and recorded using a BlackBoxToolKit USB Response Pad (Black Box ToolKit Ltd). Controls performed the task in two sessions, interleaved with a more difficult condition (data from both conditions will be reported in a separate publication). Each condition was conducted in blocks of 60 trials, with 8 blocks (4 per condition) presented per session. This setup resulted in 480 trials per session per condition, totaling 960 trials across both sessions. The order of conditions was pseudo-randomized across participants and sessions, ensuring that both conditions were equally presented within each session. PD patients performed the task as described above in one session with bilateral therapeutic stimulation switched off (DBS OFF) or with their stimulation turned on (DBS ON). EEG recordings commenced no earlier than 60 minutes after the DBS device was turned off. Each session contained 480 trials. At the end of each block, all participants received feedback on their performance using an average accuracy score. The start of each block was self-paced, allowing participants to decide when to proceed. Additionally, each session began with 30 practice trials to help participants familiarize themselves with the task. To further minimize the impact of background noise, participants wore earplugs. Stimulus presentation was carried out using MATLAB R2016b (MathWorks, Natick, USA; RRID: SCR_001622) in combination with the Psychophysics Toolbox (RRID: SCR_002881) [Brainard, 1997], running on a Windows 10 operating system.

**Figure 1:**
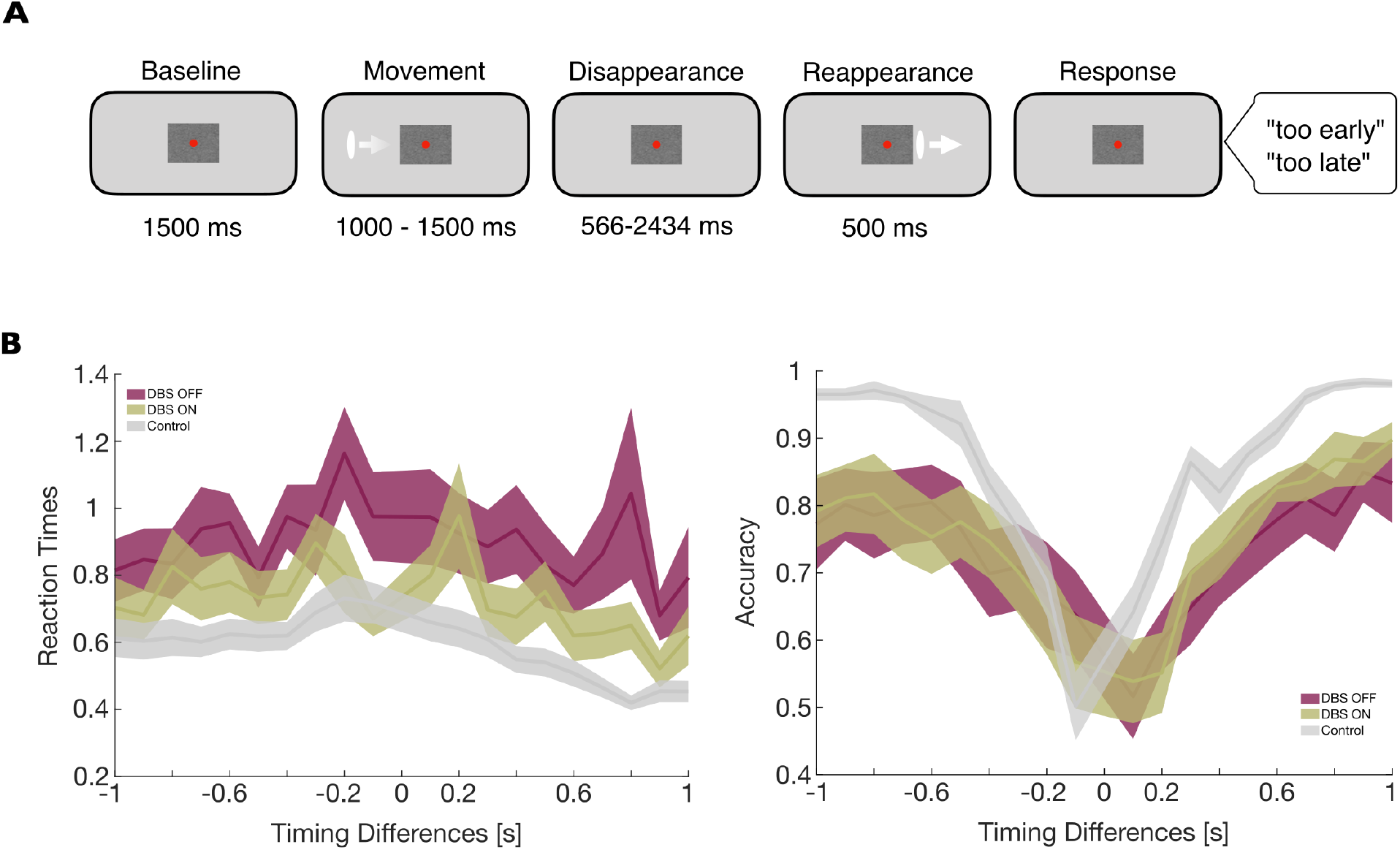
(A) Experimental design of the temporal prediction task is shown. In the beginning of each trial, a white ellipse appeared on the screen and moved towards the occluder before disappearing. After 1500 ms ± 34 to ± 934 ms the stimulus reappeared. Participants had to decide whether the stimulus reappeared *too early* or *too late* by responding with a keypress. (B) Reaction times, as well as accuracy values for all timing differences are presented as mean and shaded regions represent the standard error of the mean (SEM).

### Data acquisition and pre-processing

EEG activity was recorded using a 64-channel actiCAP snap electrode system (Brain Products GmbH, Gilching, Germany) with active Ag/AgCl electrodes embedded in an elastic cap. Each electrode was equipped with an integrated impedance converter to minimize noise from the environment and movement artifacts, which was particularly crucial for recordings in PD patients experiencing resting tremor. The data was digitized at a sampling rate of 1000 Hz using BrainAmp amplifiers (Brain Products GmbH, Gilching, Germany). Analysis was performed using MATLAB R2019b (The MathWorks Inc., 2019) and two open-source toolboxes: EEGLAB [Delorme and Makeig, 2004] (RRID:SCR_007292) and FieldTrip [Oostenveld et al., 2011] (RRID:SCR_004849), as well as the MEG and EEG Toolbox Hamburg (METH, Guido Nolte; RRID:SCR_016104) or custom-made scripts.

### EEG preprocessing/artefact removal

Initially, EEGLAB was used to extract data from the experimental blocks. Each of these blocks was filtered with a high-pass filter at 0.5 Hz to reduce slow drifts, a low-pass filter at 95 Hz to eliminate high-frequency DBS artefacts and a band-stop filter at 49.5-50.5 Hz for line noise removal. Additionally, the DBSFilt toolbox was employed to remove artefacts in the harmonics and subharmonics of the stimulation frequency. All blocks from each session were subsequently merged. The merged data were segmented into epochs of variable lengths using FieldTrip. Each trial was cut from 1240 ms before the stimulus movement onset to 1240 ms after the offset of the reappeared stimulus, resulting in trial durations ranging between 4546 ms and 6914 ms. To remove artifacts from eye movements, muscle activity, and cardiac signals, an independent component analysis (ICA) was performed using the infomax algorithm. Components were identified and rejected based on their time course, variance, power spectrum, and topography through visual inspection. On average, 19 *±* 3 out of 64 components were removed per participant per session. After ICA, all trials were visually inspected, and those containing residual artifacts not detected in earlier steps were excluded. On average, 470 *±* 5 trials per session remained after preprocessing, out of a total of 480 trials.

### Quantification and statistical analysis

Our primary focus for statistical analyses was to compare data from patients across DBS OFF and DBS ON conditions, as well as to contrast patient data with control group data. To examine differences between DBS OFF and DBS ON conditions within patients, we conducted paired-samples *t*-tests. For comparisons between the patient and control group, we utilized independent-samples *t*-tests. We accounted for multiple comparisons by adjusting the alpha level (*α*=0.025).

### Behavioral data analysis

Participants were not provided with feedback regarding the correctness of their responses. As a result, they made judgments based on their subjective interpretation of what constituted a “correct” reappearance timing. To determine these individual points of subjective equality (PSE), we fitted a psychometric curve to the behavioral data of each participant across all trials in each condition. For each timing difference, we first calculated the proportion of *too late* responses for each participant. A binomial logistic regression (psychometric curve) was then fitted to the data using the *glmfit*.*m* and *glmval*.*m* functions in MATLAB. The timing difference corresponding to a 50% *too late* response rate was identified as the point of subjective equality (PSE) for each participant. To evaluate whether there was a significant bias in the PSE values, we compared them to zero using one-sample *t*-tests. The steepness of the psychometric function was quantified as the reciprocal of the difference between the timing differences at 75% and 25% *too late* response rates, providing a measure of the slope of the curve.

### EEG analysis

We decomposed EEG recordings into time-frequency representations using complex Morlet wavelets [Cohen, 2014]. Each trial and channel were convolved with 40 wavelets logarithmically spaced between 0.5-100 Hz, with cycles increasing logarithmically from 2 to 10. Only trials labeled as correct based on individual PSEs (see 2.6) were analyzed. After the wavelet convolution, we estimated spectral power modulations by dividing trials into four overlapping windows: baseline (−550 to -50 ms), movement (−50 to 950 ms), disappearance (−350 to 950 ms), and reappearance (−350 to 450 ms), all aligned relative to their respective events. Spectral power estimates were averaged across trials, binned into 100 ms intervals, and normalized using a pre-stimulus baseline (−500 to -200 ms prior to movement onset). Grand averages across channels were tested against baseline using paired-sample *t*-tests with cluster-based permutation statistics for multiple comparison correction (cluster-*α*=0.05, 1000 randomizations). For source-space analysis, leadfields were computed using a single-shell volume conductor model [Nolte, 2003] aligned to a Montreal Neurological Institute (MNI) template with a 5003-voxel grid. Cross-spectral density (CSD) matrices were derived from wavelet-convolved data in 100 ms steps and used to compute common adaptive linear spatial filters (Dynamic Imaging of Coherent Sources (DICS) beamformer; [Gross et al., 2001]. Normalized source power was analyzed using cluster-based permutation tests to identify significant power differences between groups (two-sample *t*-tests: DBS OFF vs. Controls and DBS ON vs. Controls) and within conditions (paired-sample *t*-tests: DBS OFF vs. DBS ON). To quantify phase alignment, we computed ITPC from wavelet-convolved complex time-frequency data by extracting phase angles per trial and time point (using MATLAB-function *angle*.*m*). ITPC values (ranging from 0 to 1) were calculated for all subjectively correct trials and averaged in 100 ms bins across the four time windows (see above). Statistical comparisons between the movement, disappearance, and reappearance windows and a pre-stimulus baseline were conducted using cluster-based permutation tests for each condition and group. For source level analyses, we used the same procedure applied to spectral power (see above) and extracted ITPC estimates for each voxel. Our analyses focused on group and condition differences in the disappearance window, specifically in the delta-band (0.5-4 Hz). In all statistical comparisions, we used cluster-based permutation tests to identify significant ITPC differences between groups (independent-sample *t*-tests: DBS OFF vs. Controls and DBS ON vs. Controls) and within conditions (paired-sample *t*-tests: DBS OFF vs. DBS ON). Finally, Pearson correlations between delta ITPC and the slope of the psychometric function were computed, again, correcting for multiple comparisons through cluster-based permutation statistics.

## RESULTS

### DBS improved temporal prediction performance

To assess the effects of DBS on temporal prediction performance, we compared the slopes of the psychometric functions between and within groups. Significant differences were observed between conditions of DBS OFF and DBS ON (*t(12)=-1*.*68, p=*.*007, Cohen’s d=-1*.*7*), as well as between DBS OFF and Controls (*t(32)=-5*.*04, p<*.*001, Cohen’s d=-1*.*79*), but not between DBS ON and Controls (*t(32)=-1*.*68, p=*.*10, Cohen’s d=-0*.*60*) (Figure 2). This suggests that DBS ON improved performance in the temporal prediction task compared to DBS OFF, while performance in DBS ON was comparable to controls.

**Figure 2:**
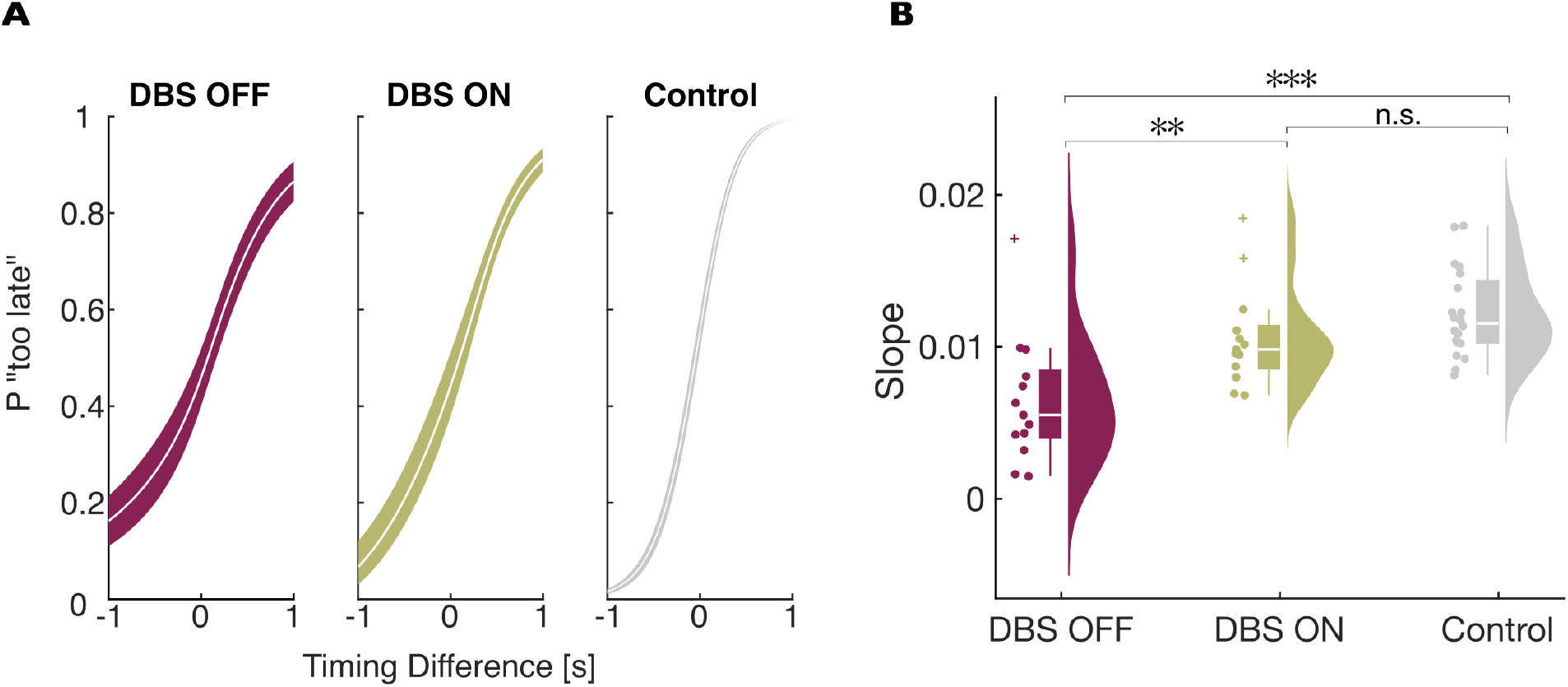
Behavioral results of the temporal prediction task. (A) Fitted psychometric curves of patients in the DBS OFF (red) and DBS ON (green) condition and healthy controls (grey). The timing difference of 0 refers to the objectively correct reappearance of the stimulus after 1500 ms. Colored areas depict standard errors of the mean (SEM) (B) Distribution of slope differences of psychometric functions between conditions and groups are shown as individual data points, boxplots and distributions. *p < .05, **p<.01, ***p<.001; n.s., non-significant.

### DBS restored cortical beta suppression during temporal prediction

The effects of DBS on task performance were associated with specific cortical activity modulation as measured by EEG. Based on previous findings [Daume et al., 2021, Gulberti et al., 2015], indicating the relevance of beta-band activity in temporal prediction, we were mostly interested in the beta frequency range after stimulus disappearance, i.e., the onset of the temporal prediction process. In controls, we observed suppression of cortical beta activity during temporal prediction compared to baseline (Figure 3A). In PD patients, this beta-band suppression was generally less pronounced in both DBS OFF and DBS ON conditions compared to controls. When observing oscillatory activity changes across DBS conditions in patients, beta power appeared to be stronger during DBS OFF than during DBS ON. This observation was supported by the results of cluster-based permutation statistics, which revealed significant clusters of voxels across bilateral medial prefrontal areas and the left temporal cortex (cluster-*p*=.019; Figure 3B). Notably, the present analysis reveals a significant decrease in beta power within the medial frontal and left temporal regions in patients with PD during the DBS OFF condition when compared to controls. However, no significant differences in beta power were observed between the DBS ON condition and the controls.

**Figure 3:**
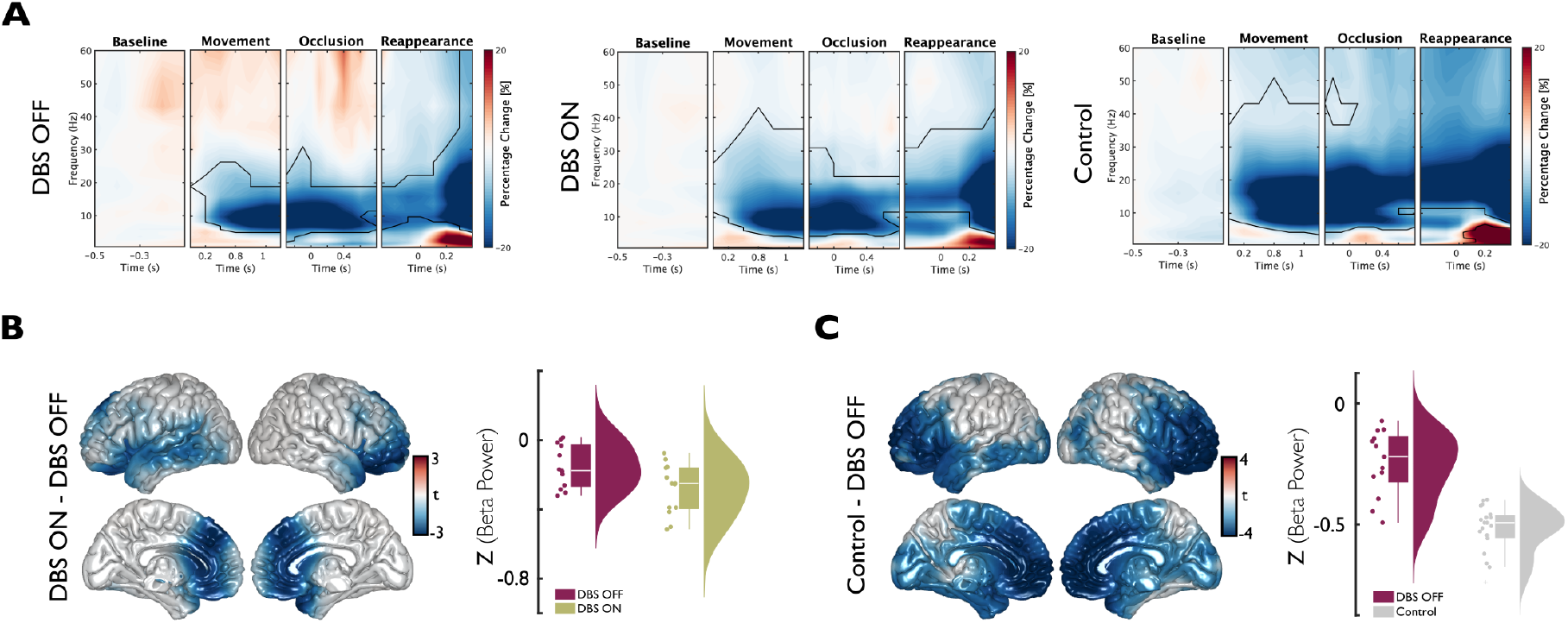
Spectral power modulations across conditions and groups. (A) Spectral power estimates compared to baseline. The panels show time-frequency plots of spectral power, averaged across sensors and participants. Each window is centered on their respective event within the paradigm and normalized to the pre-stimulus baseline. Time 0 s marks the onset of each event. Cluster-based permutation statistics revealed significant power modulations relative to baseline (indicated by regions enclosed with continuous lines). (B) Source-level comparison of beta power between DBS ON and DBS OFF conditions during the disappearance window (−0.2 to 0.6 s). Left: Surface plots of spectral beta power differences. Clusters of voxels with significant differences are highlighted in color. Right: Distribution of z-scored beta power averaged across voxels showing significant differences between conditions for the time window of disappearance. (C) Left: Source level data of beta power differences between Control and DBS OFF for the time window of disappearance (−0.2s to 0.6 s). Clusters of voxels with significant differences are highlighted in color. Right: Distribution of z-scored beta power averaged across voxels showing significant differences between groups for the time window of disappearance.

### Delta ITPC was stronger in controls compared to DBS OFF or ON

For the ITPC analysis, a similar approach to the one above was used. First, we examined ITPC differences relative to baseline within three different time windows by averaging across all sensors, using cluster-based permutation statistics. The results showed a significant increase in ITPC across various frequency ranges in time bins corresponding to movement onset, occlusion, and stimulus reappearance, respectively (all cluster-*p* <0.001; Figure 4A). The most pronounced ITPC increases for the time window centered around stimulus disappearance, i.e., the time window relevant for the temporal prediction process, were found in the delta range in patients as well as controls. Therefore, our group comparisons were focused on frequencies between 0.5 and 4 Hz in a time window of -200 to 900 ms around stimulus disappearance. When comparing delta ITPC of patients with DBS OFF or DBS ON to controls, we identified negative clusters of sensors located in frontal and occipital regions showing significant differences (all cluster-*p*<.001; Figure 4B). At source level, analysis of the -200 to 900ms time window revealed the strongest ITPC differences between patients (either DBS OFF or DBS ON condition) and controls in voxels spanning across the occipital, temporal, parietal and right frontal lobes (all cluster-*p*<0.01; Figure 4C). No significant difference of delta ITPC was found for the comparison of patients with DBS OFF vs. DBS ON condition.

**Figure 4:**
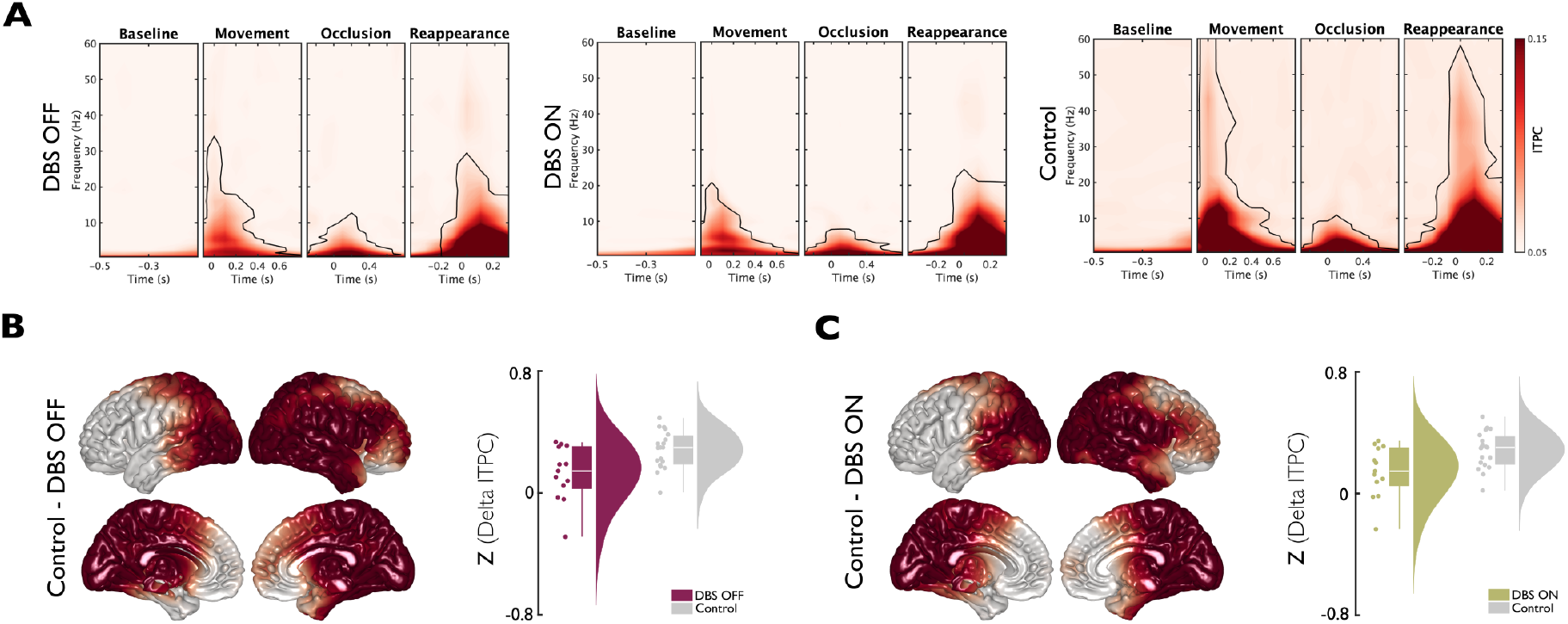
ITPC during temporal prediction in patients and controls. (A) The panels show time-frequency plots of ITPC, averaged across sensors, and participants. Each window is centered on different events within the experiment. Time 0 s marks the onset of each event. Cluster-based permutation statistics revealed significant ITPC modulations relative to baseline (indicated by regions enclosed with continuous lines). (B) Source level data of differences between Control and DBS OFF within the delta-band (0.5-4 Hz) for the time window of disappearance (−0.2 s to 0.9 s). Left: Surface plots of ITPC differences. Clusters of voxels showing significant differences between groups are highlighted in color. Right: Distribution of z-scored delta ITPC averaged across voxels showing significant differences between groups for the time window of disappearance. (C) Left: Source level comparison of ITPC data of Control and DBS ON within the delta-band (0.5-4 Hz) for the time window of disappearance (−0.2 s to 0.9 s). Clusters of voxels showing significant differences between groups are highlighted in color. Right: Distribution of z-scored delta ITPC averaged across voxels showing significant differences between groups for the time window of disappearance.

### Delta ITPC in patients with DBS ON was correlated with performance

To investigate whether the phase alignment of neural oscillations was associated with temporal predictions in patients and controls, we computed Pearson correlations of source level delta ITPC with the steepness of the psychometric function. We found a significantly positive correlation only in patients in the DBS ON condition (cluster-*p*=.012; Figure 5B). Strongest correlations were found in the cerebellum and occipital as well as parietal areas (Figure 5A). No significant correlation was found for either patients in the DBS OFF condition or controls.

**Figure 5:**
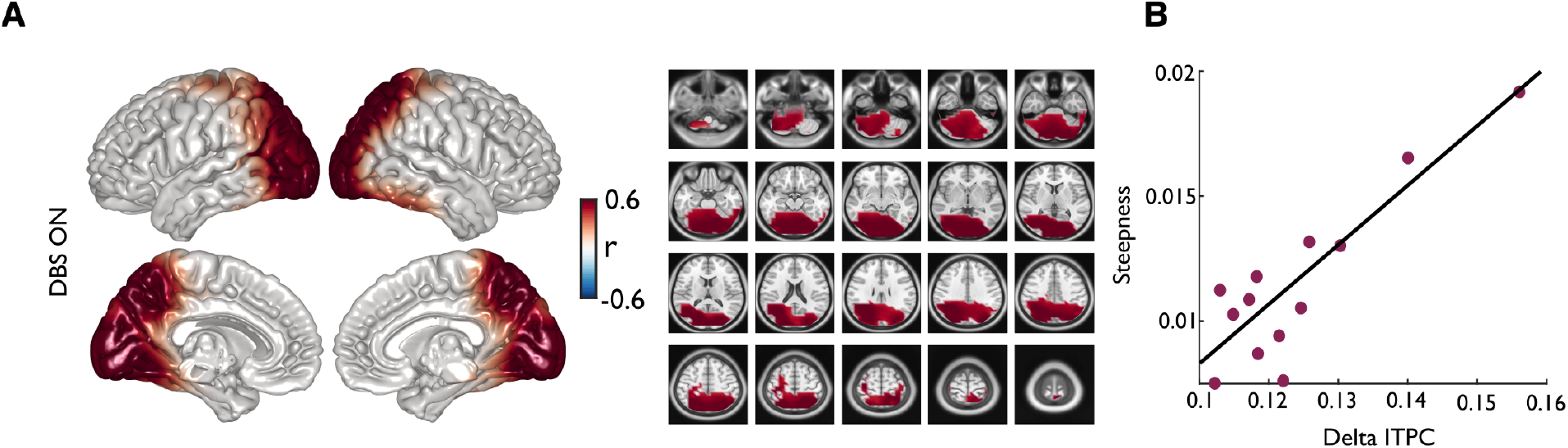
Relationship of delta ITPC and behavior. (A) Correlation of individual delta ITPC and the slope of the psychometric function within all voxels. ITPC was averaged across the delta-band (0.5-4 Hz) and time windows of -0.2 to 0.9 s around the disappearance of the stimulus. Only the clusters of voxels with significant correlations are colored. (B) Significant correlation of individual delta ITPC and steepness of the psychometric function (r=0.75, *p*=.003). Each dot of the scatter plot represents one participant. ITPC was averaged across all voxels within the clusters of significant correlations.

## DISCUSSION

The present study provides novel insights for the modulatory effects of DBS on temporal prediction in PD, offering a new perspective on how basal ganglia dysfunction impacts anticipatory cognitive processes. Consistent with our hypothesis and previous studies, PD patients showed impairments in temporal prediction relative to controls, as indicated by a shallower psychometric slope. Importantly, DBS improved performance to levels comparable to controls in the DBS ON condition. Beta suppression was less pronounced in PD patients in the DBS OFF compared to DBS ON condition and comparable between DBS ON condition and controls. In contrast, delta ITPC, was reduced in both DBS ON and OFF conditions compared to controls. For the DBS ON condition, source-level delta ITPC was positively correlated with temporal prediction accuracy. Our findings underscore that the basal ganglia play a significant role in interval timing and highlight that DBS is able to modulate higher-order cognitive functions.

### DBS and temporal prediction performance

Our behavioral results align with earlier studies showing that PD patients exhibit increased variability in timing judgments due to a slowing or disruption of the internal clock mechanism [Jones et al., 2008, Meck, 1996, Merchant et al., 2008]. These impairments have been linked to dopamine depletion in the basal ganglia, particularly the nigrostriatal pathway, which plays a central role in interval timing and temporal processing [Buhusi and Meck, 2005, Coull et al., 2011]. Within this framework, the basal ganglia are thought to modulate temporal prediction by integrating sensory and motor information across time, a process that becomes dysfunctional in PD. The restoration of temporal prediction performance in the DBS ON condition of our study suggests that STN stimulation may partially compensate for disrupted timing mechanisms. This effect may arise through the modulation of pathological oscillatory activity and partial normalization of dopaminergic transmission within the cortex-basal ganglia loop [Paulo et al., 2023]. The observed behavioral improvement of temporal prediction in our study is consistent with prior findings from studies showing that STN-DBS can enhance time perception and time reproduction tasks [Koch et al., 2004, Wojtecki et al., 2011]. Notably, our results extend these findings by demonstrating that DBS improves not only retrospective time estimation but also predictive timing, which is crucial for anticipating and responding to future events. Predictive timing is essential for a wide range of adaptive behaviors, including speech processing, motor coordination, and action planning [Arnal and Giraud, 2012, Coull et al., 2008]. Its disruption in PD may contribute not only to motor symptoms like bradykinesia but also to non-motor deficits in attention, working memory, and executive control [Allman and Meck, 2012, Buhusi and Meck, 2009]. By showing that DBS improves predictive timing, our findings suggest that the therapeutic effects of STN-DBS extend into the cognitive domain, potentially by enhancing the temporal structure of cortical processing. This has important implications for understanding the full spectrum of DBS benefits and for refining stimulation protocols aimed at cognitive enhancement in PD [Herz et al., 2018, Witt et al., 2004].

### Beta oscillations and the role of the basal ganglia

Beta oscillations have been widely associated with sensorimotor processing, typically emerging during postural stability and diminishing during active states such as movement preparation and execution [Barone and Rossiter, 2021, Kilavik et al., 2013, Pfurtscheller et al., 2003]. Furthermore, previous research has linked beta suppression to cognitive flexibility and motor readiness [Engel and Fries, 2010]. Consistent with this notion and in line with prior work [Daume et al., 2021], we observed pronounced beta power suppression in both healthy controls and patients during the temporal prediction task. However, this suppression was significantly less pronounced in patients during the DBS OFF condition, particularly in medial prefrontal and left temporal regions, compared to both patients during the DBS ON condition and healthy control participants. The reduced beta suppression in the DBS OFF condition in our study may reflect the characteristic change in cortical dynamics in PD, where excessive beta activity is associated with impaired movement initiation and reduced cognitive adaptability [Brown, 2003]. This is in line with research by Paulo et al. [2023] which highlights that cognitive impairments in PD are associated with altered beta oscillatory activity in cortex-basal ganglia circuits, specifically involving the dorsolateral prefrontal cortex and caudate nucleus. In their study, reduced beta suppression during a working memory task was associated with poorer cognitive performance in PD, suggesting a mechanistic link between beta desynchronization and cognitive function. These findings underscore the role of beta oscillations not only in motor control but also in cognitive processes, including temporal prediction. In the DBS ON condition, patients exhibited levels of beta power suppression comparable to those of healthy controls. This recovery supports the notion that STN-DBS disrupts pathological beta synchronization, thereby restoring a more flexible and responsive network state [Kühn et al., 2008, Little and Brown, 2014]. The relationship between subcortical and cortical beta is central to this dynamic. Beta activity in the STN is tightly coupled to cortical regions via the cortex-basal ganglia loop and in PD, this loop exhibits pathological synchronization and elevated beta cortico-subthalamic coherence [Brown, 2003, Lalo et al., 2008, Litvak et al., 2011, Oswal et al., 2016, Sharott et al., 2005]. Subthalamic DBS as well as administration of therapeutic doses of dopaminergic medication have been shown to attenuate beta power within the STN of PD patients [Barone and Rossiter, 2021, Kühn et al., 2009, Ray et al., 2008, Weinberger et al., 2006, Whitmer et al., 2012]. Thus, when STN-DBS reduces subcortical beta activity, it may also decouple pathological cortico-subcortical coherence [Oswal et al., 2016], enabling cortical regions, particularly motor and prefrontal areas, to re-engage in task-relevant beta desynchronization [Hemptinne et al., 2015, Whitmer et al., 2012]. The restoration of cortical beta suppression in patients during DBS ON observed in our study is consistent with this notion. Although stimulation is delivered locally to the STN, its broader therapeutic impact likely stems from the normalization of aberrant beta synchronization across interconnected cortico-subcortical pathways, thereby enabling greater functional flexibility across the system. This restoration of neural flexibility is essential for dynamic cognitive functions such as temporal anticipation, and may underlie the observed enhancements in predictive processing under DBS in individuals with PD.

### Delta-band ITPC and temporal prediction

Beyond power changes, we also examined ITPC in the delta-band, which is associated with the phase alignment of neural oscillations to expected stimuli [Arnal et al., 2015, Schroeder and Lakatos, 2009, Daume et al., 2021]. Both PD patients and healthy controls exhibited increases in delta ITPC during stimulus occlusion and reappearance, consistent with the notion that low-frequency phase alignment supports the temporal prediction of sensory input. However, both DBS OFF and DBS ON conditions in patients showed reduced delta ITPC compared to healthy controls. This might indicate that DBS did not fully restore phase alignment mechanisms. At sensor level, significant differences in delta ITPC were found in frontal and occipital regions. Source-level analyses revealed that the largest discrepancies in delta phase consistency were located in occipital, temporal, parietal, and right frontal cortices, areas involved in attentional and predictive timing [Arnal et al., 2015, Coull et al., 2008]. These findings suggest that while DBS may help recover power dynamics, it may not sufficiently restore the phase consistency necessary for optimal temporal prediction. Interestingly, the correlation between delta ITPC and psychometric function slope was only significant in the DBS ON condition, particularly in the cerebellum, occipital, and parietal regions. These are structures known to support sensorimotor timing and predictive processing [Ivry and Spencer, 2004]. This indicates that despite reduced ITPC in absolute terms, individual differences in residual phase alignment under DBS ON may still support more accurate temporal judgments. The absence of a similar relationship in DBS OFF and control groups might underscore a potentially unique, compensatory role of DBS in facilitating phase-behavior coupling.

Overall, our findings contribute to a growing literature emphasizing the role of oscillatory dynamics in cognitive processes such as temporal prediction. Specifically, our results support the view that beta and delta oscillations are key signatures of temporal prediction, and that PD disrupts both power and phase-based mechanisms. While DBS appears effective in modulating beta power and improving behavioral performance, its impact on phase-based delta coherence seems limited, warranting further investigation. Future studies could assess whether different DBS parameters (e.g., frequency, pulse width, or stimulation pattern) differentially affect beta and delta-band dynamics, respectively. For instance, applying low-frequency stimulation protocols may more effectively engage slower rhythms like delta, and could be used to causally investigate the role of low frequency phase alignment in temporal prediction.

## CONCLUSION

In summary, our study demonstrates that DBS enhances temporal prediction performance in PD by modulating key oscillatory mechanisms, particularly through beta power suppression. This is consistent with a broader body of work suggesting that beta desynchronization facilitates adaptive cognitive processes, including attention and anticipation [Engel and Fries, 2010]. However, both DBS ON and OFF conditions showed diminished delta phase alignment compared to controls, indicating that DBS may only partially restore predictive processing mechanisms. Our findings underscore the importance of integrating both power and phase-based measures when evaluating DBS effects and highlight the potential of oscillatory biomarkers to refine therapeutic strategies for cognitive dysfunction in PD. Future work combining DBS with targeted cognitive or neuromodulatory interventions may offer new avenues for restoring both the timing and precision of neural dynamics supporting temporal prediction.

## Declaration of competing interest

The authors declare that they have no known competing financial interests or personal relationships that could have appeared to influence the work reported in this paper.

### CRediT authorship contribution statement

**Rebecca Burke:** Conceptualization, Methodology, Investigation, Software, Formal analysis, Validation, Visualization, Data curation, Writing - original draft, Writing - review & editing. **Marleen J. Schönfeld**: Conceptualization, Methodology, Investigation, Formal analysis, Writing - review & editing. **Alessandro Gulberti**: Writing - review & editing. **Christian K.E. Moll**: Writing - review & editing. **Monika Pötter-Nerger**: Investigation - Assisted in patient recruitment. Writing - Review & Editing. **Andreas K. Engel**: Conceptualization, Methodology, Writing - review & editing, Funding acquisition, Project administration, Supervision.

## Acknowledgements

This work was supported by grants from the Deutsche Forschungsgemeinschaft (SFB TRR 169/B1/B4 261402652 awarded to A.K.E.). We thank Karin Reimann for assistance in data recording, and Till R. Schneider, Alexander Maÿe, and Peng Wang for helpful discussions on the data.

