## Supplementary Material for "Modulation of temporal prediction by STN-DBS in Parkinson’s disease: Links between behavior and cortical oscillations"

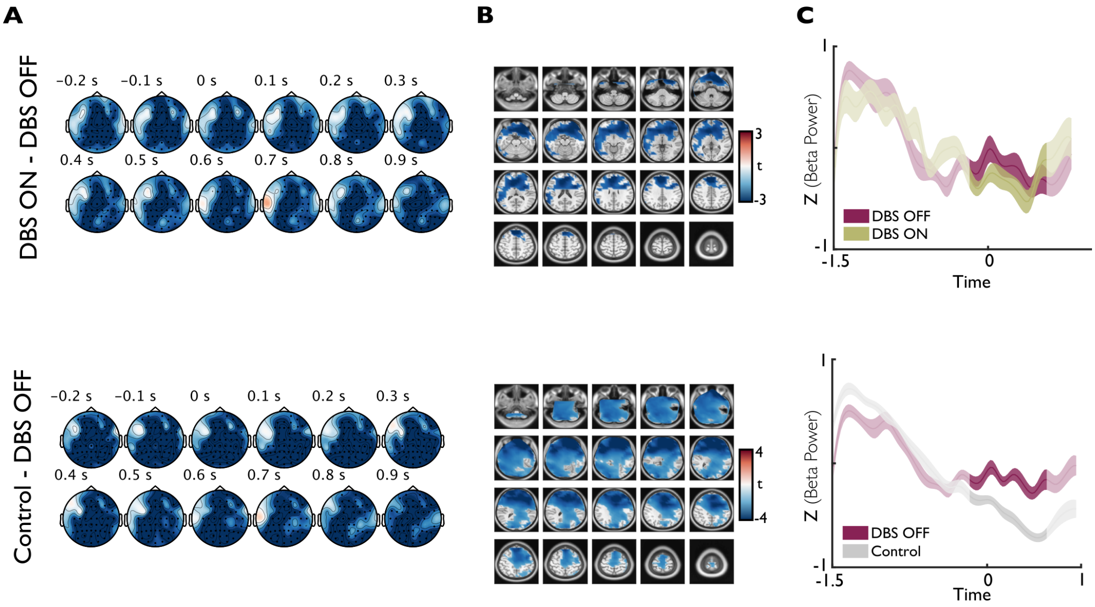


**Supplementary Figure 1.** Beta power differences between conditions (DBS ON & DBS OFF) and groups (Control & DBS OFF). (A) Sensor level data of beta power differences between DBS ON and DBS OFF (top) and Control and DBS OFF (bottom) for the time window of disappearance (-0.2s to 0.6 s). Clusters of sensors with significant differences indicated by black dots. (B) MRI slices of beta power differences between DBS ON and DBS OFF for the time window of disappearance (-0.2 s to 0.6 s). Clusters of voxels with significant differences are highlighted in color. (C) Time course of beta power (± SEM) averaged across voxels of significant clusters for the difference between conditions and groups. Time 0 s marks the onset of disappearance.

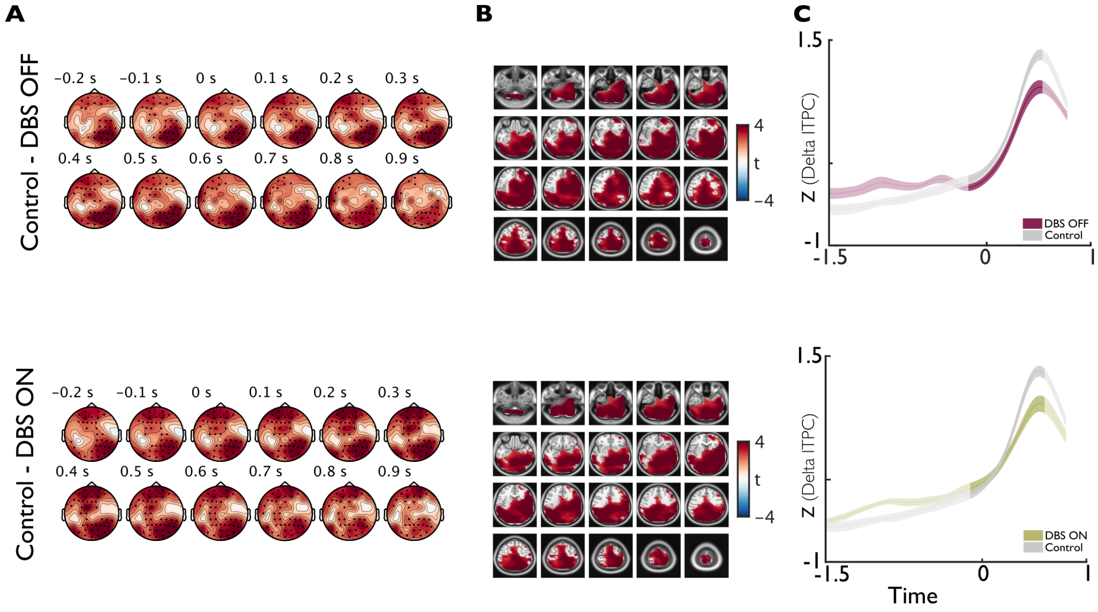


**Supplementary Figure 2.** Delta ITPC differences between groups. (A) Sensor level data of ITPC differences between Control and DBS OFF (top) and Control and DBS ON (bottom) for the time window of disappearance (-0.2s to 0.9 s). Clusters of sensors with significant differences indicated by black dots. (B) MRI slices of delta ITPC differences between DBS ON and DBS OFF for the time window of disappearance (-0.2 s to 0.9 s). Clusters of voxels with significant differences are highlighted in color. (C) Time course of ITPC (± SEM) averaged across voxels of significant clusters for the difference between groups. Time 0s marks the onset of disappearance.
